# Dissimilarity in sulcal width patterns in the cortex can be used to identify patients with schizophrenia with extreme deficits in cognitive performance

**DOI:** 10.1101/2020.02.04.932210

**Authors:** Joost Janssen, Covadonga M. Díaz-Caneja, Clara Alloza, Anouck Schippers, Lucía de Hoyos, Javier Santonja, Pedro M. Gordaliza, Elizabeth E.L. Buimer, Neeltje E.M. van Haren, Wiepke Cahn, Celso Arango, René S. Kahn, Hilleke E. Hulshoff Pol, Hugo G. Schnack

**Affiliations:** Department of Child and Adolescent Psychiatry, Institute of Psychiatry and Mental Health, Hospital General Universitario Gregorio Marañón, Madrid, Spain; Ciber del Área de Salud Mental (CIBERSAM), Instituto de Investigación Sanitaria Gregorio Marañón (IiSGM), Madrid, Spain; Department of Psychiatry, UMCU Brain Center, University Medical Center Utrecht, Utrecht, The Netherlands; School of Medicine, Universidad Complutense, Madrid, Spain; Department of Bioengineering and Aerospace Engineering, Universidad Carlos III de Madrid, Madrid, Spain; Department of Child and Adolescent Psychiatry/Psychology, Erasmus University Medical Centre, Sophia Children’s Hospital, Rotterdam, The Netherlands; Department of Psychiatry, Icahn School of Medicine at Mount Sinai, New York, United States

**Keywords:** schizophrenia, neuroimaging, morphometric similarity, cognition, longitudinal, subtyping

## Abstract

Schizophrenia is a biologically complex disorder with multiple regional deficits in cortical brain morphology. In addition, interindividual heterogeneity of cortical morphological metrics is larger in patients with schizophrenia when compared to healthy controls. Exploiting interindividual differences in severity of cortical morphological deficits in patients instead of focusing on group averages may aid in detecting biologically informed homogeneous subgroups. The Person-Based Similarity Index (PBSI) of brain morphology indexes an individual’s morphometric similarity across numerous cortical regions amongst a sample of healthy subjects. We extended the PBSI such that it indexes morphometric similarity of an independent individual (e.g., a patient) with respect to healthy control subjects. By employing a normative modeling approach on longitudinal data, we determined an individual’s degree of morphometric dissimilarity to the norm. We calculated the PBSI for sulcal width (PBSI-SW) in patients with schizophrenia and healthy control subjects (164 patients, 164 healthy controls; 656 MRI scans) and associated it with cognitive performance and cortical sulcation index. A subgroup of patients with markedly deviant PBSI-SW showed extreme deficits in cognitive performance and cortical sulcation. Progressive reduction of PBSI-SW in the schizophrenia group relative to healthy controls was driven by these deviating individuals. By explicitly leveraging interindividual differences in severity of PBSI-SW deficits, neuroimaging-driven subgrouping of patients is feasible. As such, our results pave the way for future applications of morphometric similarity indices for subtyping of clinical populations.

## Introduction

Neurobiological research and biomarker discovery efforts in the field of psychiatry have been substantially hampered by insufficient biological validity of current diagnostic categories, stalling the development of precision medicine in psychiatry (1). Given the large body of evidence for clinical, etiological and biological heterogeneity within psychotic disorders, extensive research is being conducted to identify more biologically homogeneous subgroups based on clinical, neuroimaging, neurophysiological, molecular, or biochemical variables, which might improve knowledge of the underlying pathophysiology, and guide stratified treatments (2–6).

Schizophrenia is consistently associated with gray and white matter deficits that vary in severity and some of these abnormalities are progressive over time (7–9). However, the clinical utility of brain imaging for guiding diagnosis and treatment is challenged by the large heterogeneity in location and severity of brain deficits among patients with schizophrenia (10–12). The degree of interindividual heterogeneity in brain deficits is not evenly distributed across the cortex in schizophrenia (11). Large interindividual heterogeneity in cortical deficits affecting a particular brain region in schizophrenia may be the result of separable neurobiological underpinnings of brain anatomy in that region across individuals, thus pointing to different biological subgroups within the disorder. In contrast, regional brain deficits with low interindividual variability may point to mechanisms shared by a significant proportion of patients with schizophrenia, thus supporting their involvement in its general pathophysiology (11–13). Collectively, these findings underline the importance of focusing on interindividual differences in severity in addition to assessing mean differences (14).

Novel morphometric approaches assessing the similarity of cortical morphology across regions, instead of focusing on single regions, may thus be well suited for assessing the widespread, variable cortical deficits in schizophrenia (15,16). A recent cross-sectional study showed that in three independent samples of adults with schizophrenia, average global and regional morphometric similarity was reduced, and this reduction was associated with schizophrenia-related genes (15). However, it is unclear if reductions in morphometric similarity in schizophrenia are progressive over time. In addition, reduced morphometric similarity was based on mean differences, which, in combination with widespread enlarged dispersion of cortex morphology in schizophrenia, suggests that results may not be equally applicable to all patients. The recently developed Person-Based Similarity Index (PBSI) combines the concepts of morphometric similarity and interindividual heterogeneity by calculating an individual’s morphometric similarity to the other individuals in a group across cortical regions (16).

To identify patients whose brain morphology is markedly dissimilar to that of a normative group, we extended the PBSI for patients such that it indexes the degree of similarity between the morphological profile of an individual patient to those of healthy control subjects. We chose to use sulcal width as morphological measure, thus calculating PBSI for sulcal width (PBSI-SW). Increased sulcal width has been related to schizophrenia (17) and to decreases in cortical thickness, and reductions in cortical gray and white matter, thus suggesting that changes in sulcal width may reflect the result of a combined effect of gray and white matter atrophy (18–20). Sulcal morphology assessments are particularly suited for lifespan studies, as they are based on MRI gray matter-cerebrospinal fluid contrast. Measurements based on gray-white matter contrast (e.g. cortical thickness) might be confounded by the age-dependency of this contrast when using standard MRI sequences (21).

We leverage a large longitudinal lifespan sample of participants with schizophrenia and healthy participants to investigate whether the lifespan PBSI-SW trajectories differ between patients and healthy subjects, and whether such differences are driven by individuals with extremely deviating PBSI-SW. In addition, we investigate whether extreme PBSI-SW deviance is associated with deficits in cognition and -in line with the neurodevelopmental hypothesis of schizophrenia-the sulcation index, a marker of perinatal neurodevelopmental perturbations and which is associated with schizophrenia (22,23).

## Methods and Materials

### Sample

From a large longitudinal sample of patients with schizophrenia and healthy participants aged 16-70 years (at baseline) we included individuals who had T1-weighted magnetic resonance imaging (MRI) scan acquisitions at two time points. Detailed information regarding diagnostic criteria and clinical assessments of the Utrecht Schizophrenia project and the Genetic Risk and Outcome of Psychosis (GROUP) consortium, Utrecht, The Netherlands are described in (22–24). In brief, patients with a DSM-IV diagnosis of schizophrenia were recruited in various inpatient and outpatient facilities. A sample of healthy controls was recruited in the same geographic areas. Participants in the patient and control groups with major medical or neurological conditions, or an estimated intelligence quotient (IQ) below 80 were excluded. Further details on the specific inclusion and exclusion criteria for each cohort can be found in the Supplement. The IRB at the University Medical Center Utrecht reviewed the study protocols and provided ethical approval. All participants provided written informed consent.

For purposes of the current study, we included images of patients who fulfilled DSM-IV criteria for schizophrenia both at baseline and follow-up (Figure 1A), had sufficient scan quality (QA procedures are described in more detail in the Supplement), and matched patient and control groups for sex, resulting in 164 healthy participants and 164 patients, contributing a total of 656 scans. Demographic, cognitive, clinical and imaging information for the final sample can be found in Table 1. For every participant, we recorded age at scan, sex, handedness, and estimated IQ values (based on four subtests of the short forms of the Wechsler Adult Intelligence Scale (WAIS) or the WAIS III, see (25) for further details). For the patient group, clinical severity was assessed using Positive and Negative Syndrome Scale (PANSS) scores at baseline and at follow-up (26). Duration of illness was calculated by subtracting age at the onset of illness from age at the time of scan.

**Table 1.** Demographics, cognitive, clinical and imaging characteristics of patients with schizophrenia and healthy controls. The total amount of scans is 912. ^1^Age of onset is the age at which the first positive symptom occurs. ^2^Duration of illness is calculated based on the date of the appearance of first positive symptoms of schizophrenia until the date of scan. ^3^Antipsychotic treatment data is not available for all the patients included (82 patients had medication information). Cumulative doses were calculated per time of scan and given in chlorpromazine equivalents using standard conversion factors and estimated by the daily doses of each of the antipsychotics used by the patient. Significance calculated by Chi-square or Welch t-tests when appropriate. Significant diagnostic differences are for one or both timepoints. *<0.05, **<0.01, ***<0.001. IQ, Intelligence quotient; PANSS, Positive and Negative Symptom Scale.

|  | Schizophrenia patients |  | Healthy controls |  | P |
| --- | --- | --- | --- | --- | --- |
|  | Baseline | Follow-up | Baseline | Follow-up |  |
| Number of subjects | 164 |  | 164 |  |  |
| Sociodemographics |  |  |  |  |  |
| Sex, n (%), males | 126 (77) |  | 126 (77) |  |  |
| Age, years: mean (SD) | 29.52 (8.95) | 33.60 (9.21) | 31.04 (11.65) | 35.05 (11.88) |  |
| Education, years: mean (SD) | 11.94 (2.68) | 12.11 (2.85) | 13.61 (2.65) | 13.66 (2.63) |  |
| Cognitive and clinical variables |  |  |  |  |  |
| Estimated-scale IQ total: mean (SD) | 97.62 (16.56) | 102.97 (20.47) | 112.59 (15.78) | 115.46 (16.11) | *** |
| Age of onset, years: mean (SD)<sup>1</sup> | 21.63 (5.70) |  |  |  |  |
| Duration of illness, years: mean, (SD)<sup>2</sup> | 6.77 (7.61) | 11.23 (8.27) |  |  |  |
| PANSS score, total: mean (SD) | 62.78 (18.78) | 50.24 (14.30) |  |  |  |
| Total Positive symptoms | 14.83 (5.78) | 12.36 (4.63) |  |  |  |
| Total Negative symptoms | 16.17 (6.01) | 12.09 (5.66) |  |  |  |
| Total General symptoms | 30.88 (11.04) | 24.99 (7.05) |  |  |  |
| Antipsychotics<sup>3</sup> |  |  |  |  |  |
| Without medication, n (%) | 7 (4.14) | 7 (4.16) |  |  |  |
| Exclusively typical, n (%) | 23 (13.6) | 12 (7.14) |  |  |  |
| Exclusively atypical, n (%) | 89 (52.66) | 92 (54.76) |  |  |  |
| Both, n (%) | 4 (2.36) | 2 (1.19) |  |  |  |
| Cumulative dose exposure (mg):<br>mean (SD) | 275837.8 (397345.9) | 945754.9 (1285851) |  |  |  |
| Missing, n (%) | 103 (61.54) | 102 (60.71) |  |  |  |
| Imaging |  |  |  |  |  |
| Total brain volume | 1181.27(112.70) | 1171.50 (115.42) | 1218.15 (111.20) | 1213.57 (110.68) | ** |

**Figure 1.**
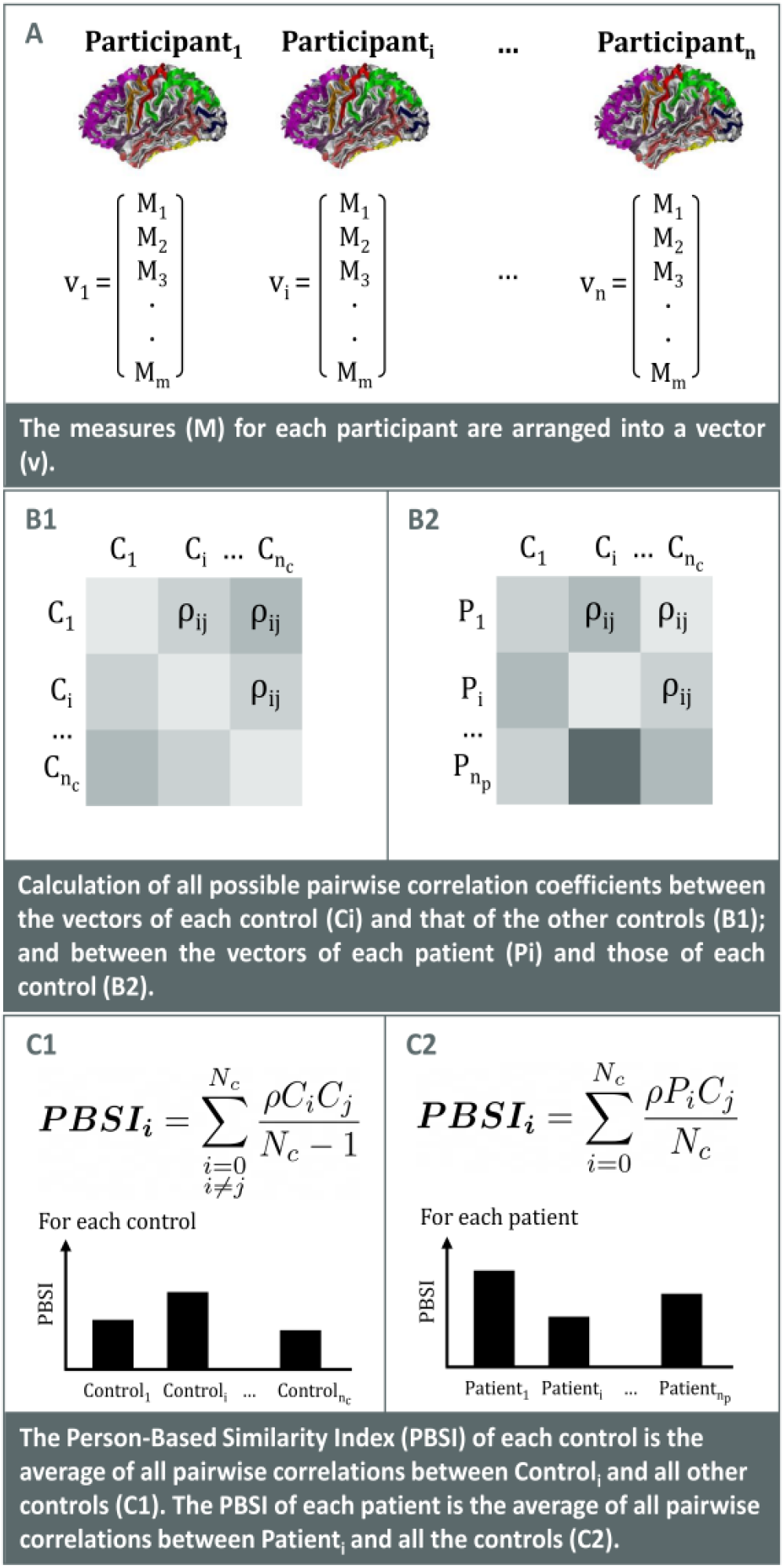
Pipeline for computing a Person-Based Similarity Index (PBSI) for healthy controls (based on (16)) and patients.

Data on antipsychotics usage around time of scan was retrieved: we determined the cumulative intake (until around the time of each scan) of antipsychotic medication converted into chlorpromazine milligram equivalents (CPZ) per participant.

#### MRI acquisition and image analysis

Two scanners (same vendor, field strength and acquisition protocol) were used. Participants were scanned twice on either a Philips Intera or Achieva 1.5 T and a T1-weighted, 3-dimensional, fast-field echo scan with 160-180 1.2 mm contiguous coronal slices (echo time [TE], 4.6 ms; repetition time [TR], 30 ms; flip angle, 30°; field of view [FOV], 256 mm; in-plane voxel size, 1×1 mm^2^) was acquired. All included participants had their baseline and follow-up scan on the same scanner.

#### Image processing

Images were analyzed using the FreeSurfer analysis suite (v5.1) with default settings to provide detailed anatomical information customized for each individual (27–29). The FreeSurfer analysis stream includes intensity bias field removal, skull stripping, and generation of a “ribbon” image and reconstruction of gray and white matter surfaces. Total brain tissue volume was derived as the sum of total gray and white matter volumes. For all images, sulcal segmentation and identification was performed with BrainVISA software (v4.5) using the Morphologist Toolbox using default settings (30). Sulcal width and the sulcation index were calculated using the default protocols within BrainVISA (17,31). Using BrainVISA sulci nomenclature the recognized sulci were pooled in eleven (a priori determined) bilateral areas, identical to the regions used by (32). See the Supplement for a detailed description of BrainVISA image processing, the calculation of sulcal width and the sulcation index, BrainVISA nomenclature and a graphical illustration of the eleven regions. FreeSurfer and BrainVISA derived measurements have been validated via histological and manual measurements and have demonstrated to show good test–retest reliability (18,33–35).

### PBSI-SW

The PBSI-SW calculation was based on (16). Briefly, at each time point and for each individual we computed a PBSI-SW value using a five-step procedure (see Figure 1). First, we created the sulcal width profile of each participant by concatenating the corresponding morphometric measures (sulcal width for eleven bilateral sulcal regions). Second, for each of the *n_c_* controls we calculated the interindividual Spearman’s rank correlation coefficients between the participant’s sulcal width profile and the profiles of the other *n_c_*-1 controls. Third, for each patient we calculated the interindividual Spearman’s rank correlation coefficient between the participant’s sulcal width profile and the profiles of all *n_c_* controls. Fourth, for each control the *n_c_*−1 interindividual correlation coefficients for sulcal width were averaged to yield one PBSI-SW per control. Fifth, for each patient the *n_p_* interindividual correlation coefficients for sulcal width were averaged to yield one PBSI-SW per patient. Higher PBSI-SW (with a maximum of 1) denotes greater similarity between an individual’s sulcal width profile and those of the (other) controls. The R script used for the PBSI-SW calculation is available from github.

### Statistical analyses

All analyses were performed in R (https://cran.rstudio.com/).

#### Longitudinal trajectory of PBSI-SW

To investigate the longitudinal trajectories of PBSI-SW over the age range, generalized additive mixed models (GAMMs) were used (36). The GAMM fitting technique represents a flexible routine that allows nonparametric fitting with relaxed assumptions about the relationship between brain morphological metrics and age (37). The technique is well suited for fitting nonlinear relationships through local smoothing effects, independent of any predefined model, and robust to age-range selections and distant data points (38). GAMM models were implemented to examine age, diagnosis (i.e., patient vs control), as well as an age×diagnosis interaction while controlling for scanner, total brain volume and the random effect of the individual. We first fitted models including a sex and age×sex term but these terms were not significant, we therefore excluded these terms from the model. Including scanner as a random effect did not change the results. To better understand the age×diagnosis interaction, GAMM estimates for age were also implemented and visualized for patients and controls separately.

#### Cross-sectional comparisons

To compare PBSI-SW between groups at baseline and follow-up separately, PBSI-SW values were residualized for age, scanner, and total brain volume, at each time point and within each diagnostic group. Next, after adding back group mean values for each metric we used the Welch t-test to assess differences between the diagnostic groups at baseline and follow-up.

#### Normative modeling

First, we created a normative reference for PBSI-SW, by calculating the average and standard deviation of the residualized PBSI-SW values for the healthy control subjects. Then, individual deviance of a participant’s PBSI-SW with respect to the normative group was determined as follows. We calculated for each participant at each time point the distance between his/her residualized PBSI-SW value and the normative residualized PBSI-SW value in terms of standard deviations, i.e. a PBSI-SW-Z value for each participant. This PBSI-SW-Z value reflects how deviant the PBSI-SW value of a given individual is compared to the normative value at a certain time point, independently of the participant’s age, scanner and total brain volume. For comparison, we also calculated Z-values using the method described in (39). The Pearson correlation coefficient between them was 0.94 (see SFigure 3). Then, we applied a threshold, |PBSI-SW-Z| > 2, (as in (40)) to identify individuals whose morphometric profile is markedly dissimilar (at any time point) to the morphometric profiles of the normative group: the deviants.

To assess the impact of deviance on the PBSI-SW lifespan trajectories we recalculated GAMM models without healthy control and patient deviants. To investigate whether the group of patients with deviant PBSI-SW shared differential clinical, cognitive and morphological characteristics relative to the rest of the patients we compared age at baseline, sex, PANSS total scores and positive, negative and general subscores at baseline, estimated IQ at baseline, and sulcation index at baseline and follow-up between the deviant and non-deviant patient groups. Finally, we also assessed whether image quality differed in patients at the extremes of the distributions.

#### Regional contribution to PBSI-SW

To calculate the regional contribution to the PBSI-SW values we used a leave-one-region-out method, i.e., recalculating each individual’s PBSI-SW after removing one region at a time. The region’s contribution was calculated by subtracting the recalculated residualized PBSI-SW values from the original residualized PBSI-SW value (as in (16)). Average regional contributions were calculated for healthy controls, non-deviant patients and deviant patients at baseline and follow-up. A larger positive contribution meant that the recalculated PBSI-SW decreased more, i.e., the region contributed more to the original PBSI-SW value. A more negative contribution value meant that excluding the region increased the recalculated PBSI-SW more, i.e., the region had a larger negative impact on the original PBSI-SW.

The regions were then ranked by contribution value from healthy control participants at baseline. For each region, contributions were compared between healthy controls, patient non-deviant and deviant groups using the Welch t-test and corrected for multiple comparisons using FDR (baseline and follow-up, 66 comparisons). Effect sizes for mean differences are given as Cohen’s d.

## Results

### Longitudinal trajectory of PBSI-SW

The PBSI-SW trajectories were significantly different between groups (age-by-diagnosis interaction: F=14.57, p<0.001, see Table 2 and Figure 2A). Patients displayed a close to linear reduction with age. Controls showed an increase of PBSI-SW until 30 years approximately after which PBSI-SW remained stable. Cross-sectional comparisons of PBSI-SW between patients and controls at baseline and at follow-up demonstrated a significant difference with small to intermediate effect size at follow-up (mean PBSI-SW for controls = 0.80, for patients = 0.78, t = 3.23, df = 278.16, p = 0.001, mean difference 0.02, 95% confidence interval (CI): 0.01,0.04, Cohen’s d = 0.36), see Figure 2B.

**Table 2.** Generalized Additive Model estimates for age, diagnosis, scanner, total brain volume and age*diagnosis for Person Based Similarity Index for Sulcal Width (PBSI-SW) before and after outlier removal. Smooth function (edf) as well as degrees of freedom (Ref.df) and F-statistic and associated significance (***=p<0.001;**=p<0.01;*=p<0.05).

| **Intercept** | **Estimate** | **SE** | **t** | **p** |
| --- | --- | --- | --- | --- |
| Diagnosis | -1.302e-02 | 7.032e-03 | -1.851 | 0.0649 |
| Total brain volume | 6.446e-05 | 3.100e-05 | 2.079 | 0.0382 |
| Scanner | 7.521e-03 | 7.407e-03 | 1.015 | 0.3105 |
| **Slope** | **edf** | **Ref.df** | **F** | **p** |
| s(age) | 2.204 | 3 | 11.864 | 0.07815 |
| s(age):patients | 1.042 | 3 | 14.570 | 0.00975 |
| Outliers removed |  |  |  |  |
| **Intercept** | **Estimate** | **SE** | **t** | **p** |
| Diagnosis | -7.168e-03 | 3.941e-03 | -1.819 | 0.069694 |
| Total brain volume | 5.944e-05 | 1.781e-05 | 3.338 | 0.000919 |
| Scanner | 1.031e-03 | 4.172e-03 | 0.247 | 0.804988 |
| **Slope** | **edf** | **Ref.df** | **F** | **P** |
| s(age) | 2.5759 | 3 | 24.974 | 0.000234 |
| s(age):patients | 0.5552 | 3 | 1.217 | 0.139211 |

**Figure 2:**
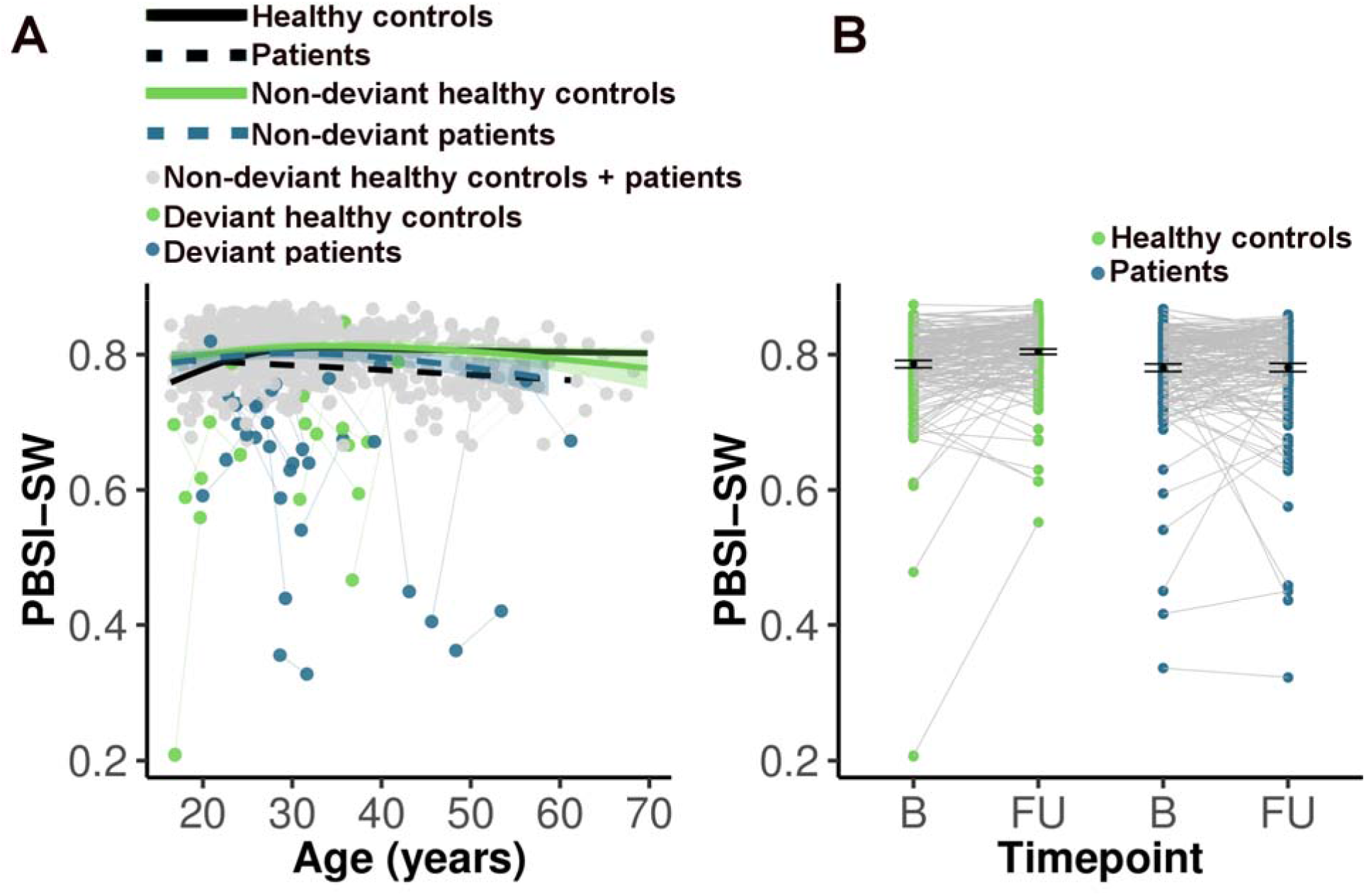
**A**) Generalized additive mixed models (GAMM) for Person Based Similarity Index for Sulcal Width (PBSI-SW). Four fits over the spaghetti plot are shown: 1) healthy controls, 2) patients, 3) non-deviant healthy controls, 4) non-deviant patients. Deviants were participants with |PBSI-SW-Z| > 2, i.e., markedly deviating from the normative morphometric value at any time point. GAMMs included total brain volume and scanner as covariates. The age×diagnosis interaction was no longer significant after removal of the deviants. **B**) PBSI-SW values for healthy controls and patients at baseline and follow-up. PBSI-SW was residualized for age, scanner and total brain volume. B, baseline; FU, follow-up.

### PBSI-SW deviance (PBSI-SW-Z)

As expected, more patients than controls had significant deviance at any of the time points: 10 controls had PBSI-SW-Z < −2 and 19 patients (χ^2^ = 3.06, df = 1, p = 0.04). Of these 19 patients, 16 patients showed a decrease in PBSI-SW-Z between baseline and follow-up (see Figure 3A), which is, assuming equal proportions of increase and decrease, a disproportional high number (χ^2^ = 8.89, df = 1, p < 0.01). In non-deviant patients the effect was opposite, 57 out of 145 participants showed a decrease (χ^2^ = 6.63, df = 1, p = 0.01). We recalculated lifespan PBSI-SW trajectories excluding deviants and the age×diagnosis interaction was no longer significant (see Table 2 and Figure 2A)

**Figure 3:**
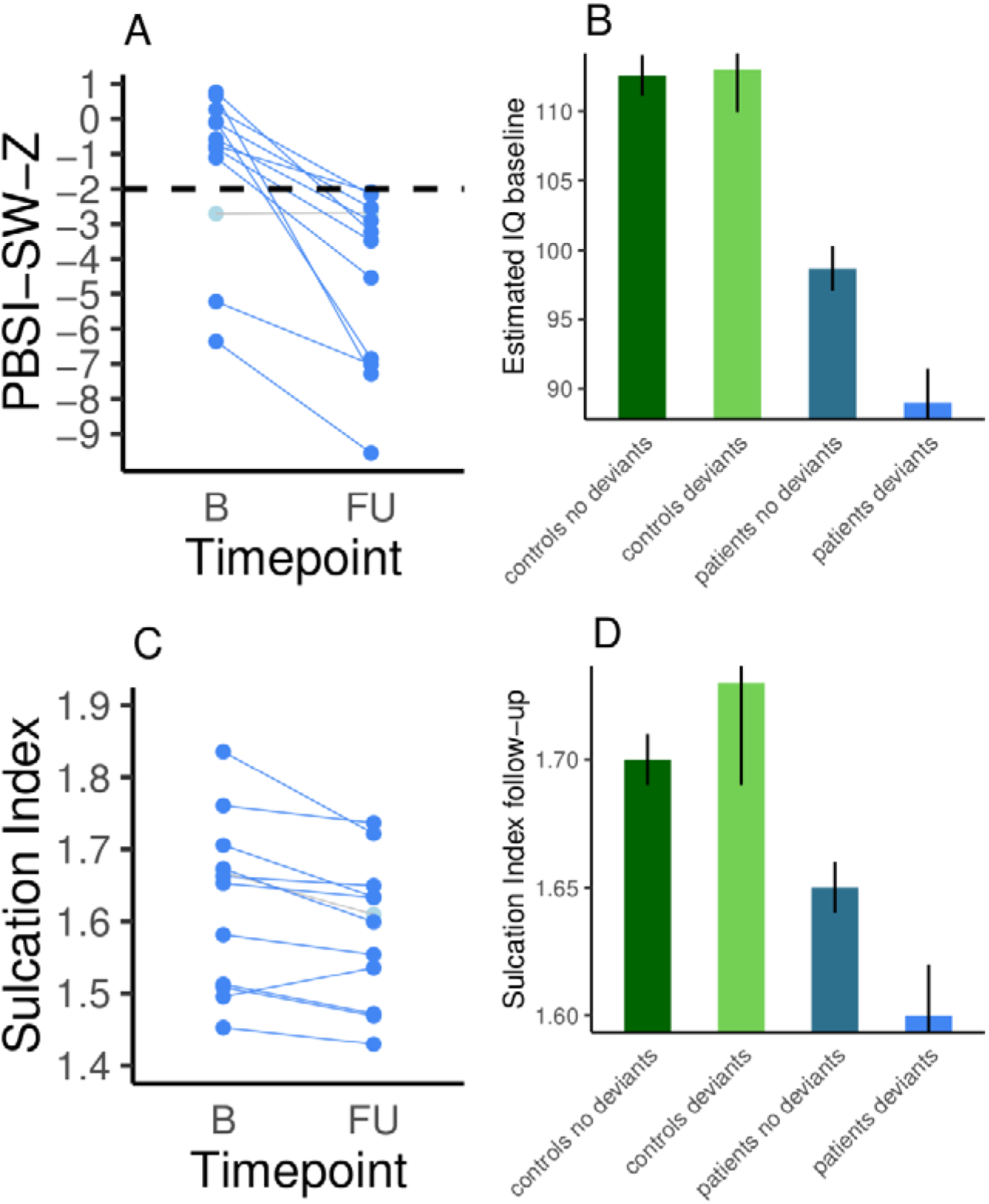
**A**) Person Based Similarity Index for Sulcal Width Z values (PBSI-SW-Z) for thirteen deviant patients (six deviant patients did not have estimated IQ at baseline available) with PBSI-SW-Z < −2 at any time point. All but one patient had a more negative PBSI-SW-Z value at follow-up as compared to baseline. **B**) Average and standard error for estimated Intelligence Quotient (IQ) at baseline for controls and patients, non-deviant patients and deviant patients. **C**) Sulcation Index at baseline and follow-up for deviant patients. **D**) Average sulcation index and standard error at follow-up for controls and patients, non-deviant patients and deviant patients.

The PBSI-SW-Z < −2 deviant group of 19 patients did not differ from the other patients in terms of age, sex, scanner, baseline total PANSS scores, positive, negative and general PANSS subscores, or image quality metrics (see SFigure 4), but they did differ in estimated IQ at baseline (mean (se) estimated IQ non-deviants: 98.66 (1.64), deviants: 89.00 (2.14), t = −3.32, df = 21.06, p < 0.01, mean difference: −9.66, 95%CI: − 16.46,−3.79, Cohen’s d = −0.62, see Figure 3B). In order to deal with potential problems related to unbalanced and small sample sizes we repeated the comparison of estimated IQ between the deviant patient group and the non-deviant group using non-parametric permutation testing (1000 permutations, F = 4.145, p < 0.05).

**Figure 4:**
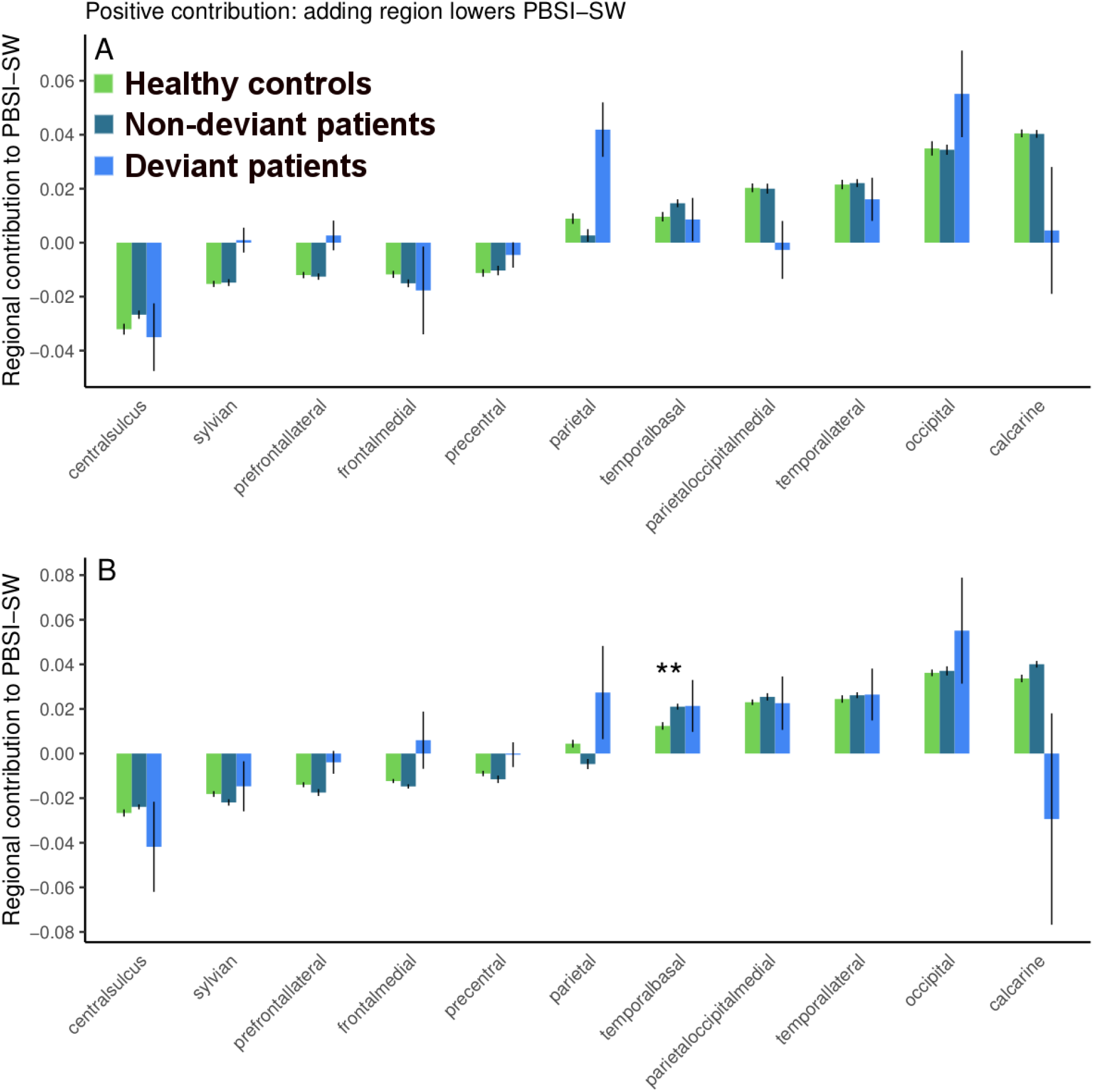
Average and standard error of each region’s contribution value to the Person Based Similarity Index for Sulcal Width (PBSI-SW) for healthy controls, non-deviant and deviant patients (PBSI-SW-Z < −2 at any time point) at baseline (**A**) and at follow-up (**B**). Regional contribution values to PBSI-SW were calculated as (average PBSI-SW(all-regions-included) - average PBSI-SW(leave-one-region-out)), thus a negative contribution value means that leaving the region out increased the PBSI-SW, i.e., the region had a negative effect on the PBSI-SW. Regions are ranked by contribution values for healthy controls at baseline. **: p<0.01 between controls and non-deviant patients after FDR correction (baseline and follow-up, controls vs non-deviant patients, controls vs deviant patients, non-deviant patients vs deviant patients).

Patients had a lower sulcation index compared to healthy control subjects at follow-up (mean (se) sulcation index healthy control subjects: 1.68 (0.01), patients: 1.64 (0.01), t = −3.44, df = 321.79, p < 0.01, mean difference −0.04, 95%CI: −0.06,−0.02, Cohen’s d = −0.38). The deviant patient group had a lower sulcation index compared to the non-deviant patient group (mean (se) sulcation index non-deviants: 1.65 (0.01), deviants: 1.60 (0.02), t = −2.40, df = 13.25, p = 0.03, mean difference −0.05, 95%CI: − 0.13,−0.01, Cohen’s d = −0.76, see Figure 3C and 3D below). For patients, estimated IQ at baseline and sulcation index at baseline and follow-up were not significantly associated with PBSI-SW (all p-values > 0.05, see Figure S5)

In controls, the PBSI-SW-Z < −2 deviant group did not differ from the non-deviant healthy control group in estimated IQ at baseline (mean (se) estimated IQ non-deviants: 112.57 (1.49), deviants 113.00 (3.08), t = 0.13, df = 10.63, p = 0.90, mean difference 0.43, 95%CI: −7.12,7.99, d = 0.03) and sulcation index at baseline (mean (se) sulcation index non-deviants: 1.68 (0.01), deviants: 1.71 (0.05), t = 0.72, df = 9.52, p = 0.49, mean difference 0.03, 95%CI: −0.08,0.15, d = 0.34) or follow-up (mean (se) sulcation index non-deviants: 1.70 (0.01), deviants: 1.73 (0.04), t = 0.59, df = 9.65, p-value = 0.57, mean difference 0.03, 95%CI: −0.07,0.13, d = 0.25).

### Regional contribution to PBSI

In controls, regional contribution showed a similar pattern at baseline and follow-up with frontal regions contributing negatively to PBSI-SW (see Figure 4 and Figure 5), i.e. leaving the region out increased the PBSI-SW. This pattern was also present in non-deviant patients. For the temporal basal region at follow-up the contribution was significantly higher in non-deviant patients compared to controls after correction for multiple comparisons (t = −4.032, df = 270.77, p < 0.01, mean difference −0.01, 95%CI: − 0.013,−0.004, Cohen’s d = −0.45). The deviant patient group did not differ from the other groups in contribution value for any of the regions.

**Figure 5:**
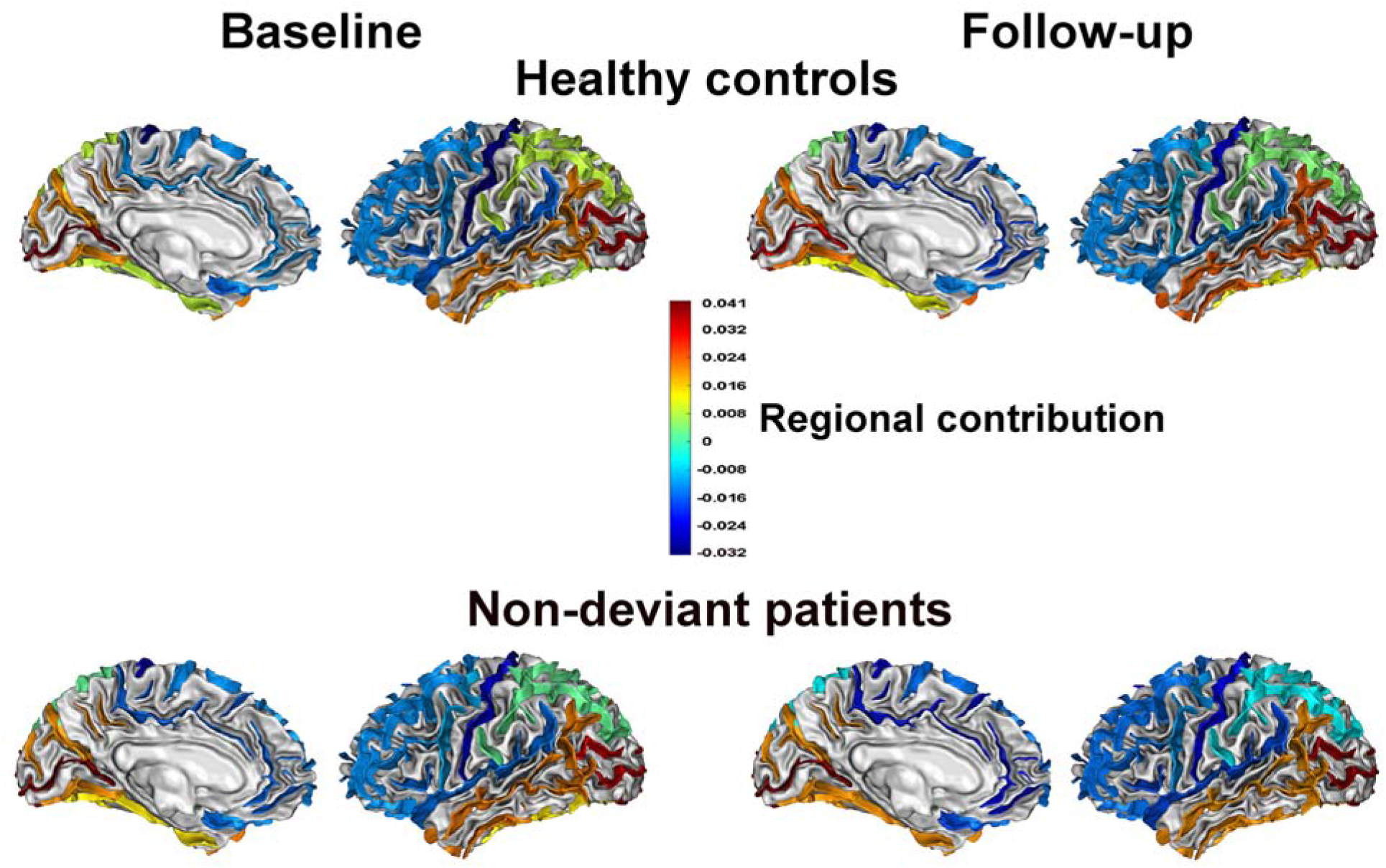
Baseline and follow-up heat maps for the contribution of each of the 11 sulcal regions to the Person Based Similarity Index for Sulcal Width (PBSI-SW) in healthy controls and non-deviant patients. Sulci are identified by sulcal median meshes (30). The regional contribution value to PBSI-SW was calculated as (average PBSI-SW(all-regions-included) - average PBSI-SW(leave-one-region-out)), thus a negative contribution value implies that leaving the region out increases the PBSI-SW, i.e., the region has a negative effect on the PBSI-SW.

## Discussion

This study is the first to use the Person Based Similarity Index (PBSI), a recently developed metric that quantifies variation in brain structural profiles across the cortex at the level of the individual. We extended the PBSI such that it quantifies the similarity between the sulcal width (SW) profile of an individual patient with schizophrenia to that of healthy control subjects. This approach allowed us to index for each individual patient the level of deviance of PBSI of sulcal width (PBSI-SW) with respect to a normative group (i.e., healthy control subjects). Our main finding is that significant deviance of PBSI-SW was present in a small group of patients only. These patients had more severe deficits in estimated IQ, and a lower sulcation index at follow-up when compared to non-deviating patients and controls. On average, schizophrenia was associated with progressive reduction of PBSI-SW over the lifespan when compared to controls, but this diagnostic effect was primarily driven by the relatively small subgroup of participants who deviated markedly in PBSI-SW.

Schizophrenia is characterized by complex and heterogeneous neurobiological and genetic underpinnings. Indeed, schizophrenia is a highly polygenic condition, with many common alleles of small effect size cumulatively conferring risk for the disorder (41). Brain structural deficits are spread out over the cortex and there is great variability in the pattern of regional deficits among patients (7,12). This makes global measures such as the PBSI-SW suitable metrics as they summarize deviations in multiple regions instead of focusing on a single region (16). Using the adapted PBSI approach we were able to translate the heterogeneity of the pattern of sulcal width deficits present in patients with schizophrenia into variation in a single number, PBSI-SW. However, in contrast to the traditional case-control approach, this variation could be used as a measure of individual deviance and therefore facilitate detection of biologically more homogeneous subgroups with schizophrenia (14).

Marked deviance of PBSI-SW was sparse, limited to only a small subset of patients. This finding is in line with recent reports using normative modeling showing that deviance (with respect to a control group) for cortical thickness was present only in small subsets of patients with schizophrenia, bipolar disorder or autism spectrum disorders (39,40,42). Furthermore, in patients with bipolar disorder, greater deviance was associated with worse performance on tasks of processing speed and executive functioning but not with age. However, deviance was not used to classify patients into subgroups (39). The current study extends these findings by demonstrating that while PBSI-SW did not correlate significantly with cognitive performance and global sulcation in the whole group, patients with markedly deviating PBSI-SW had lower cognitive performance and decreased global sulcation compared to non-deviating patients. The clinical importance of PBSI-SW deviance was further underlined by our finding that the effect of diagnosis on change of PBSI-SW over time was driven by the same small subset of deviating patients. This stresses further the limitations of focusing on common effects in traditional case-control designs to appropriately appraise the complex neurobiology of psychiatric disorders.

Leveraging the heterogeneity among patients with schizophrenia, bipolar disorder or schizo-affective disorder, patients with one of these three psychotic disorders have been regrouped into three biotypes using brain electrophysiological and neuropsychological measurements; these subgroups were validated by assessing the patients’ structural brain deficits (2). The biotype that clustered on low cognitive control performance also had the severest structural brain deficits compared to the other biotypes but diagnostic categories were spread out over the three biotypes. These findings together with ours demonstrate large interindividual differences in severity, type and location of structural brain deficits in schizophrenia and strongly suggest that criteria from diagnostic manuals do not adhere to neurobiology. As such, proposed alternative approaches such as the Research Domain Criteria may be important for harmonizing clinical characterization and neurobiological underpinnings (1).

Abnormally increased sulcal width and decreased sulcation index have been associated with schizophrenia, bipolar disorder and senescence (17,20,21,31). Schizophrenia has been associated with aberrant early life neurodevelopmental processes. Indeed, exposure to adverse environmental factors during fetal life may increase the risk of developing psychotic disorders (43). Sulcal morphology is strongly linked to early life neurodevelopmental processes responsible for changing the cortical surface from lissencephalic to its archetypical folded appearance (18,44). The sulcation index may be used to retrospectively assess potential impairments in these processes. However, the sulcation index also reduces during adolescence as a consequence of cortical thinning and white matter growth (19). Synaptic pruning, trophic glial and vascular changes and/or cell shrinkage in combination with genetics may be underlying decreases in sulcation (18,45,46); some of these processes may be particularly pronounced in schizophrenia (47,48).

Although we did not find differences between deviants and non-deviants on imaging quality metrics, we cannot rule out subtle effects (e.g. more severely ill patients moving slightly more during scan acquisition and taking more medication) on the results. Future studies focusing on this question are warranted. IQ was estimated from the performances on four subtests of the Wechsler Adult Intelligence Scale III; this procedure may show some limitations in particular populations relative to using full scale IQ scores (49). However, it has shown good validity as a measure of general cognitive ability, thus supporting the association between marked deviance and poorer cognitive performance (50). Although we assessed a large sample of patients with schizophrenia, the small proportion of patients with considerable deviance in PBSI-SW led to small sample sizes for the clinical and cognitive characterization. If applied to larger (multicenter) samples, our methodological approach could enable identification of larger groups with marked deviance to further characterize this phenotype using additional clinical and cognitive measures. Moreover, although the sample was well characterized, with diagnostic, clinical and cognitive assessments conducted by experienced professionals, we did not have information about development or premorbid adjustment. Adding such variables, and extending the range of age at onset with patients with adolescent-onset or even childhood-onset schizophrenia, could aid in characterizing the subgroup with marked deviance, especially considering its association with deficits in sulcation index, as a measure of developmental impairments.

## Acknowledgements

This work was supported by the Spanish Ministry of Science, Innovation and Universities. Instituto de Salud Carlos III (PI16/02012, PI17/01249, PI17/00997. PI19/01024), co-financed by ERDF funds from the European Commission, “A way of making Europe”, CIBERSAM. Madrid Regional Government (B2017/BMD-3740 AGES-CM-2), European Union Structural Funds. European Union Seventh Framework Program under grant agreements FP7-HEALTH-2009-2.2.1-2-241909 (Project EU-GEI), FP7-HEALTH-2009-2.2.1-3-242114 (Project OPTiMISE), FP7-HEALTH-2013-2.2.1-2-603196 (Project PSYSCAN) and FP7-HEALTH-2013-2.2.1-2-602478 (Project METSY); and European Union H2020 Program under the Innovative Medicines Initiative 2 Joint Undertaking (grant agreement No 115916, Project PRISM, and grant agreement No 777394, Project AIMS-2-TRIALS), Fundación Familia Alonso, Fundación Alicia Koplowitz and Fundación Mutua Madrileña. Dr. Díaz-Caneja holds a Juan Rodés grant from Instituto de Salud Carlos III (JR19/00024).

The authors thank Zimbo Boudewijns, Joyce van Baaren and Diego Beltrán for technical assistance.

