## Supplement for "Dissimilarity in sulcal width patterns in the cortex can be used to identify patients with schizophrenia with extreme deficits in cognitive performance"

**Table of contents**

Sample selection and assessment	3

Image quality assessment	5

**SFigure 1**: Flowchart summarizing the exclusion criteria process	7

Sulcal Width and Sulcation Index	9

**STable 1**: Sulcal regions with BrainVISA nomenclature	11

**SFigure 2**: Eleven sulcal regions	13

Comparison with Nispat	14

**SFigure 3**: Correlation of PBSI-SW-Z with PBSI-SW-Z Nispat	14

Image quality metrics	15

**SFigure 4**: Image quality metrics of deviant vs non-deviant patients	15

Estimated IQ and Sulcation Index	16

**SFigure 5**: Correlation PBSI-SW with estimated IQ and Sulcation Index	16

References	17

**Sample selection and assessment**

Participants for the current study were selected from a large longitudinal sample based on two cohorts including patients with schizophrenia and healthy controls, the Utrecht Schizophrenia project and the Genetic Risk and Outcome of Psychosis (GROUP) consortium, in Utrecht, the Netherlands. In both cohorts, patients with schizophrenia recruited in various inpatient and outpatient facilities and an age-matched sample of controls recruited in the same catchment areas were included. Participants with major medical or neurological illnesses or with an estimated IQ below 80 were excluded.

In the Utrecht Schizophrenia project, the Comprehensive Assessment of Symptoms and History (CASH) and the Schedule for Affective Disorders and Schizophrenia Lifetime version assessed by trained psychiatrists and psychologists were used for diagnostic assessment (1,2). Patients had to fulfill DSM-IV diagnostic criteria for schizophrenia or schizophreniform disorder at baseline, and had a 1-year follow-up confirmed diagnosis of schizophrenia. Healthy controls met Research Diagnostic Criteria of “never being [mentally] ill”. Participants in the patient and control samples were aged 16-70 years, and both samples were matched for age and sex (3). Exclusion criteria for both samples were the following: i) IQ below 80, ii) major medical or neurological illness (including migraine, epilepsy, hypertension, cerebrovascular disease, cardiac disease, diabetes, or endocrine disorders), iii) past head trauma, iv) alcohol or other substance dependence.

For the GROUP consortium, diagnostic assessments were conducted by trained raters with the CASH at baseline and the Schedules for Clinical Assessment in Neuropsychiatry (SCAN) at follow-up (2,4). Patients had to fulfill DSM-IV diagnostic criteria for a non-affective psychotic disorder at baseline and a diagnosis of schizophrenia or schizoaffective disorder at follow-up (5,6). For healthy control participants inclusion criteria were absence of a lifetime diagnosis of psychotic disorder and absence of a positive family history of a psychotic disorder in first- or second- degree relatives, as established with the Family Interview for Genetic Studies (7). Participants in the patient and control samples were aged 16-50 at baseline.

**Image quality assessment**

After the completion of the preprocessing pipeline for all T1-weighted MRI scans in FreeSurfer, we rigorously assessed the quality of the data using a combination of visual inspection and the examination of several quantitative quality assessments (QA) metrics. First, we calculated five QA measurements based on the ones proposed by the Preprocessed Connectome Project (<http://preprocessed-connectomes-project.org/quality-assessment-protocol/#spatial-anatomical>) to identify images that were unusable: signal-to-noise ratio (SNR), contrast to noise ratio (CNR), Foreground to background Energy Ratio (FBER), percent artefact Voxels (Artifacts) and Entropy Focus Criterion (Entropy). Following ENIGMA criteria (<http://enigma.ini.usc.edu/protocols/imaging-protocols/>) we defined the threshold for outliers as [mean-(2.698*SD)] for SNR, CNR and FBER metrics; and Artifacts and Entropy were tested for [mean+(2.698*SD)]. Next, for each scan the whole brain mean cortical thickness, total cortical surface area, total white matter volume, total gray matter volume, subcortical gray matter volume and intracranial volume were calculated. Then, we summed for each image the amount of outliers over all measures and calculated the mean+(2.698*SD) for the amount of outliers over the whole sample and designate those above the threshold as outliers.

QA-based exclusions

After the evaluation of the computed quantitative measures of image quality, 138 scans were considered of insufficient quality and were excluded from our analysis (see SFigure 1A). These excluded scans were validated by manually assessing image quality after preprocessing to assure that all preprocessing steps worked and to avoid the dissemination of error along the analysis. The parameters that were most useful to objectively detect artefacts visually were agreed on between researchers, e.g. incorrect sulcal labelling or insufficient quality of sulcal segmentation resulting in gross anatomical abnormalities. These visual checks were performed in BrainVisa. Those scans with poor initial sulcal labelling were manually edited, which often resulted in better sulcal labelling when manually inspected (see SFigure 1B) for examples of “good”, “medium” and “bad” quality sulcal labelling after first preprocessing of the images). In addition, 14 scans were excluded after this manual inspection.

**SFigure 1**. **A**) Flowchart summarizing the exclusion criteria process: from an initial sample of 1824 total scans, a total of 760 scans were excluded for subsequent analysis. The final sample before matching for sex is comprised of 912 scans: a. indicates the number of included scans; b. the number of excluded scans and c. the exclusion criteria (initial and imaging criteria) applied to the sample. **B**) Illustrates examples of sulcal segmentation of different qualities performed with BrainVisa: good quality of sulcal reconstruction, medium quality of sulcal reconstruction, included after manual sulcal editing; and bad quality scan which shows a lump caused by incorrect sulcal reconstruction which caused exclusion. **C**) Graphical representation of sulcal width and the sulcation index (SI). Left: sulcal width is defined as the distance between each gyral bank averaged over all points along the median sulcal surface. Right: total hemispheric sulcal surface is defined as the sum of the area of all sulcal median meshes (blue), brain hull area (transparent triangulated surface) is derived by calculating the area of the brain mesh defined via a morphologic closing of the brain mask which excludes sulcal areas during definition of the exposed cortical surface area (8). *SI*, sulcation index; *A_sulcus_*, the hemispheric sulcus surface area; *A_brain hull_^hemisphere^* , the brain hull area.

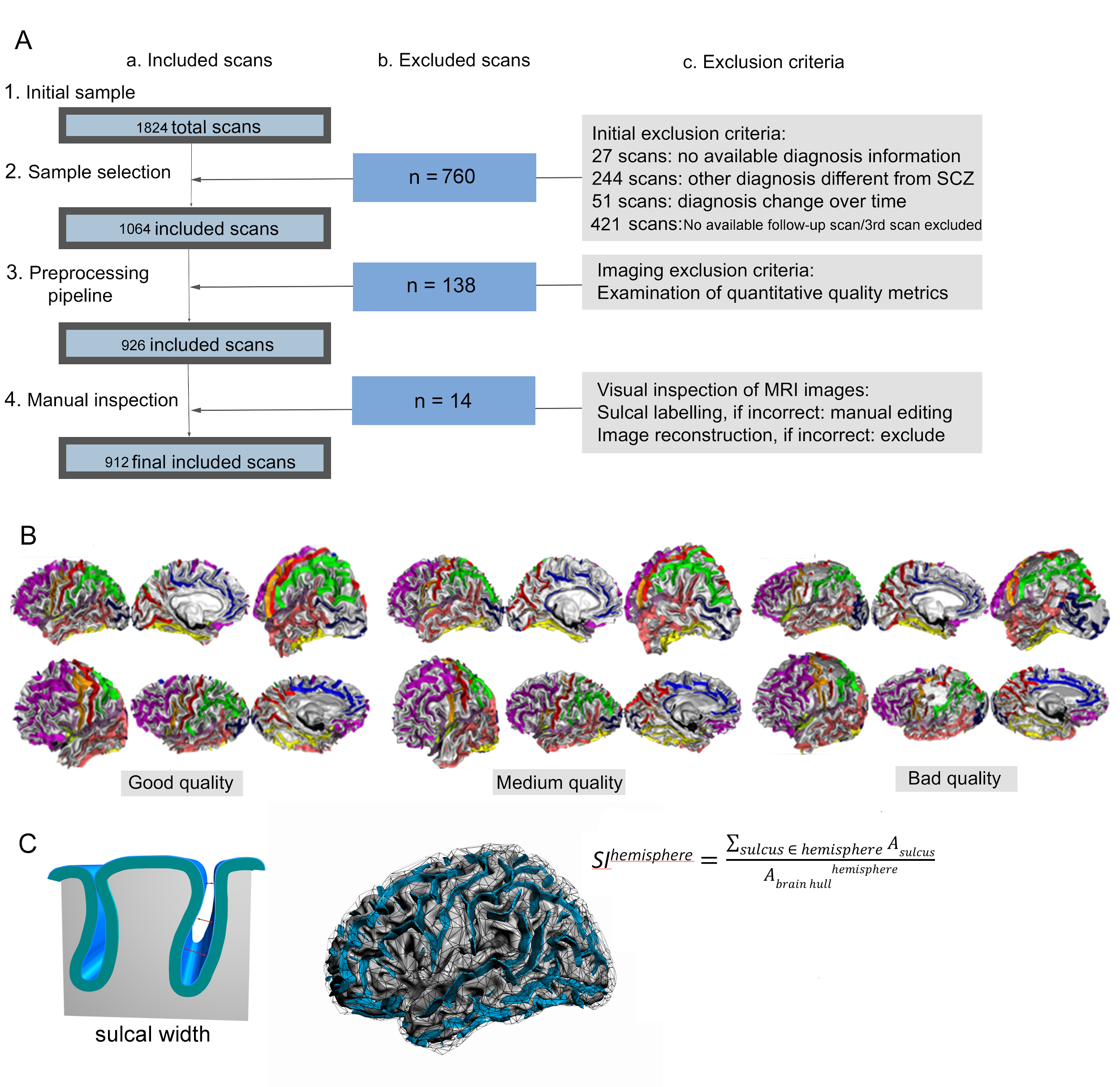

**Sulcal Width and Sulcation Index**

After importing the “ribbon” image into BrainVISA each sulcus is segmented with the cortical sulci corresponding to the crevasse bottoms of the “landscape,” the altitude of which is defined by image intensity. The brain hull area is derived in native space by calculating the area of the brain mesh defined via a morphologic closing of the brain mask that ensures boundary smoothness, which excludes sulcal areas during definition of the exposed cortical surface area. The median sulcal surface spans the entire space contained in a sulcus, from the fundus to its intersection with the brain hull. The median sulcal surface areas are summed to provide a measure of total sulcal surface area for each participant. The sulcation index (SI) is the ratio between total sulcal surface area and brain hull area:

1)
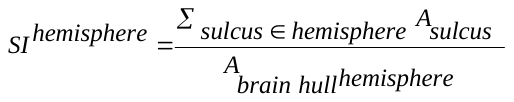

with *A_sulcus_*, the hemispheric sulcus surface area and *A_brain hull_^hemisphere^* , the brain hull area. A cortex with extensive folding has a large SI, whereas a cortex with low degree of folding has a small SI. At a constant outer cortex area, the SI increases with the number and area of sulcal folds, whereas the SI of a lissencephalic cortex is zero. SI describes the burying of the cortex and is therefore slightly different from the classical gyrification index, the ratio of the whole gyral contour length to the outer, exposed surface, which embodies additional information (included in the whole gyral contour length) related to the cortex thickness and sulcal width. Sulcal width is defined as the distance between each gyral bank averaged over all points along the median sulcal surface, see SFigure 1C for a graphic representation of sulcal width and the sulcation index. Using BrainVisa sulci nomenclature the recognized sulci were pooled in eleven (a priori determined) bilateral areas, identical to the regions used by (32). A list of which sulci were pooled to which region can be found in STable 1 and their visualization in SFigure 2. Sulcal width and the sulcation index were measured in the native space of the participant's images and left and right hemisphere values were averaged.

**STable 1**. Regions with their respective sulci

The regions are based on the eleven regions from (9). Names of the sulci follow the BrainVisa nomenclature which can be found at: http://brainvisa.info/web/_static/images/bsa/nomenclature.png.

| Region | Sulci |
| --- | --- |
| Calcarine | F.Cal.ant.-Sc.Cal. |
| Frontal Medial | F.C.M.ant.; F.C.M.post.; F.C.M.r.AMS.ant.; S.Call.; F.C.M.sup.r.asc.ant.; S.p.C.; S.C.LPC.; S.F.int.AMS.; S.F.int.; S.F.int.pol.; S.F.int.sup.; S.R.sup.; S.R.inf. |
| Parietal Occipital Medial | F.P.O.; S.s.P.; S.Pa.int.; S.Pa.t.; S.Li.; S.Li.ant.; S.Li.post.; S.Cu. |
| Sylvian Fissure | F.C.L.p.; F.C.L.r.sc.post.; F.C.L.a.; F.C.L.r.ant.; F.C.L.r.asc.; F.C.L.r.diag.; F.C.L.r.sc.ant. |
| Occipital | OCCIPITAL.; S.O.p. |
| Parietal | F.I.P.Po.C.inf.; F.I.P.Po.C.sup.; F.C.L.r.retroC.tr.; S.Pa.sup.; F.I.P.Horiz.; F.I.P.ParO.; F.I.P.r.trans.; F.I.P.r.int.1.; F.I.P.r.int.2.; S.Po.C.sup.; S.GSM. |
| Prefrontal lateral | S.F.median.; S.F.polaire.tr.; S.F.marginal.; S.F.orbitaire.; S.Or.; S.Or.l.; S.Olf.; S.F.sup.; S.F.sup.post.; S.F.sup.ant.; S.F.sup.moy.; S.F.inter.; S.F.inter.ant.; S.F.intern.moy.; S.F.inter.post.; S.F.inf.; S.F.inf.ant.; S.F.inf.moy.; S.F.inf.post. |
| Central Sulcus | S.C.; S.C.sup.; S.C.inf.; S.C.sylvian.; |
| Precentral | S.Pe.C.sup.; S.Pe.C.marginal.; S.Pe.C.median.; S.Pe.C.inf.; S.Pe.C.inter. |
| Temporal basal | S.O.T.lat.ant.; F.Coll.; S.Rh.; S.O.T.lat.post.; S.O.T.lat.int.; S.O.T.lat.med. |
| Temporal lateral | S.T.s.; S.T.pol.; S.T.i.post.; S.T.i.ant.; S.T.s.ter.asc.post.; S.T.s.ter.asc.ant. |

**SFigure 2**. The eleven sulcal regions used in the study.

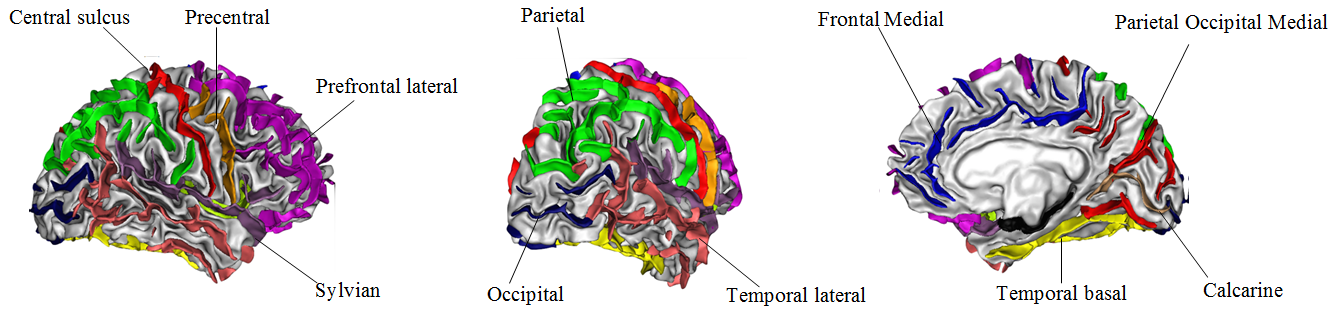

Lateral, posterior and medial view of the eleven sulcal regions (as in(9)) used in the current study.

**SFigure 3**. Pearson correlation between Z-values for sulcal width from the current study with those from Nispat (10). r=Pearson correlation coefficient; CI, confidence interval.

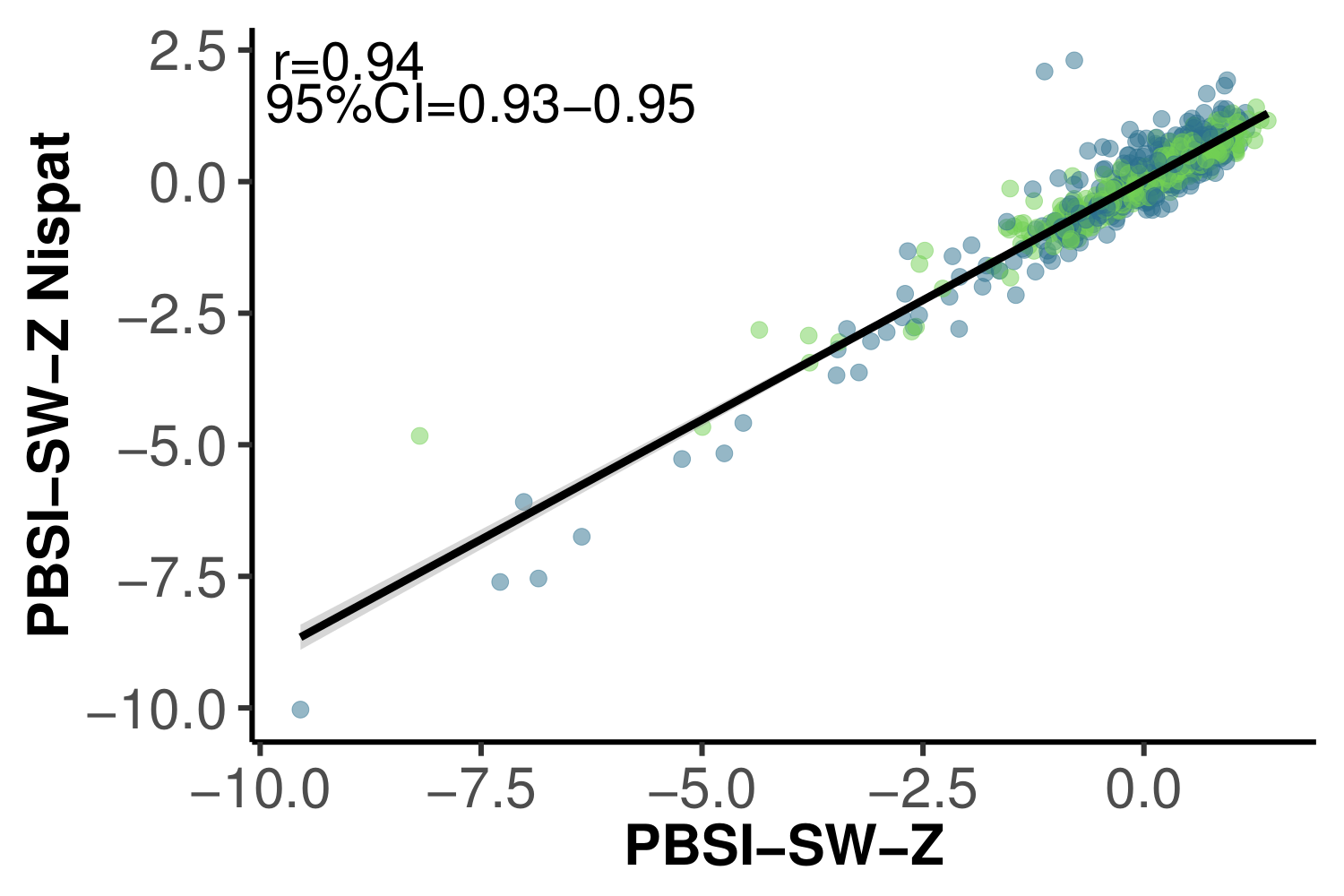

**SFigure 4**. Means and standard errors for the non-deviant and deviant patients for five image quality metrics. SNR, Signal-to-Noise Ratio; CNR, Contrast-to-Noise Ratio; FBER, Foreground to Background Energy Ratio; Artifacts, percent artifact voxels; Entropy, Entropy focus criterion. For more information about the image quality metrics see <http://preprocessed-connectomes-project.org/quality-assessment-protocol/#spatial-anatomical>.

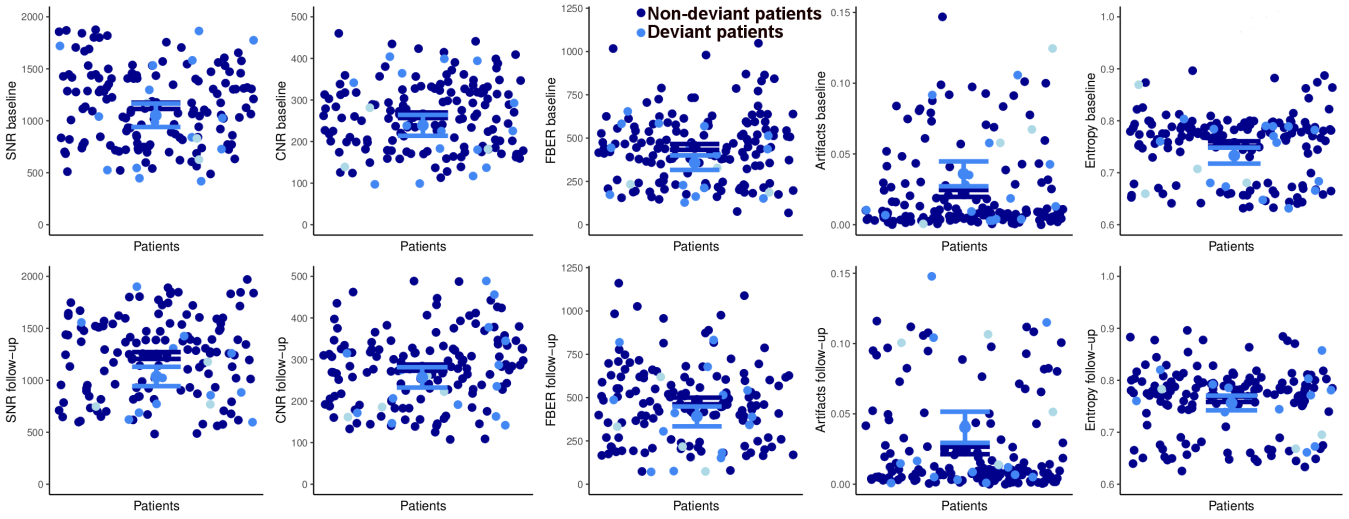

**SFigure 5**. Plots showing the non-significant (all p’s > 0.05) association between **A**) estimated Intelligence Quotient (IQ) at baseline and Person-Based Similarity Index for sulcal width (PBSI-SW) and **B**) sulcation index and PBSI-SW in patients. Solid line represents the linear association at baseline.

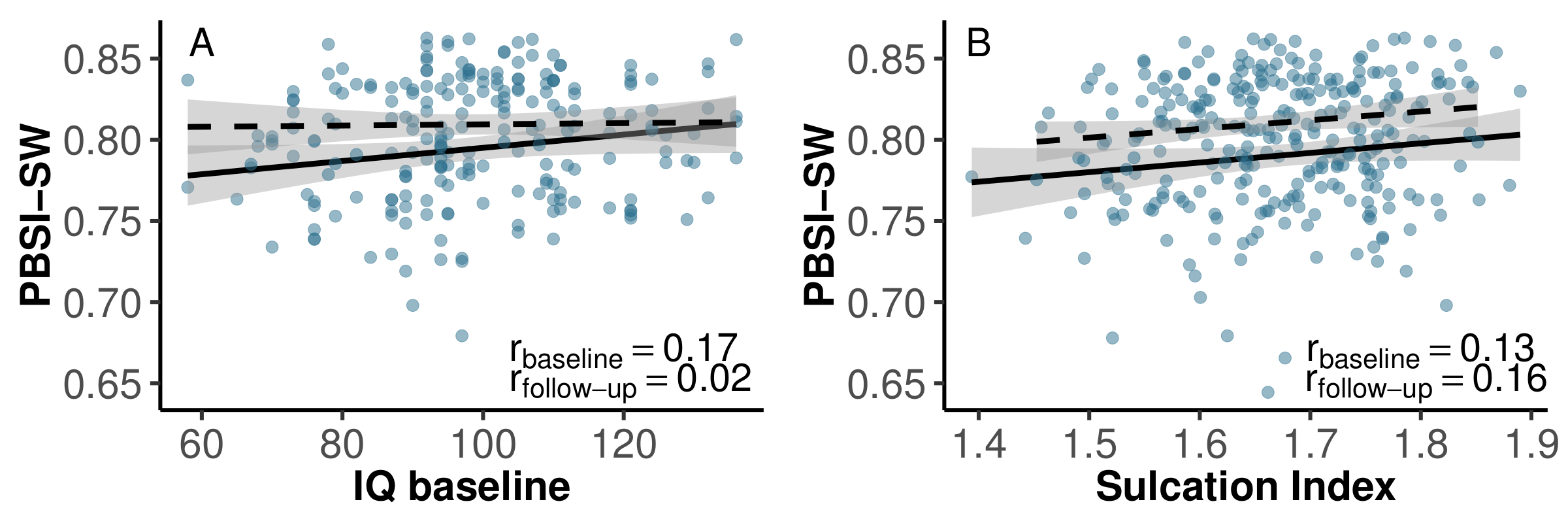
